# “Magnetic Sand”: Illusions of Interactivity

**DOI:** 10.1101/2024.07.03.598775

**Authors:** Shinsuke Shimojo, Kensuke Shimojo, Eiko Shimojo, Shengjie Zheng, Daw-An Wu

## Abstract

We found a series of new illusions, in which actions performed near a random white-noise display lead to the perception that the display is altered interactively with the observer’s actions. The perceptions resemble interactions with a box of magnetic sand, where the hand can leave traces, or attract and repulse grains in its vicinity. 1) the observer puts a finger very close to a dynamic noise display, slowly moving as though drawing a letter or a shape. A trace appears left in the finger’s path, decaying within 500 ms or so. 2) When the observer moves their palm toward and away from the display, opening and closing their fingers as if grabbing and releasing grains of sand, the random dots appear as though they were magnetically attracted to or repelled by the fingers. 3) When an open hand close to the display is slowly moved back and forth laterally, the nearby dots appear to get attracted to or captured by the fingers and thus appear to move with them. 4) The same kind of action capture occurs even when the hand is not visible, moving behind the display. These illusions are robust across a wide range of parameters, including frame rate, luminance contrast, dot size (spatial frequency), and finger movement factors. Inter-subject variability is not correlated across illusion types, and the illusions also diverge in behavior across dynamic and static noise conditions. This indicates that multiple mechanisms are involved to different extents across illusions. Several known visual motion detectors and other low-level mechanisms may be involved in seeding the perceptual phenomena. However, a complete explanation would require mechanisms of action capture, whereby the internal model of the person’s actions and their predicted consequences organizes visual attention and processing of the random stimulus components.

## 1 Introduction

Visual motion ambiguity leaves freedom for neural processing to resolve the motion perception in multiple ways. Carefully controlled visual ambiguity can thus reveal underlying processing mechanisms, causing illusions that visually embody the computations involved. New illusions reported with such ambiguous stimuli[1–3] have turned out to be especially informative in scientists’ efforts to understand the neural mechanisms underlying visual motion perception. Here, we employ a dynamic visual white noise display (fully ambiguous motion) to reveal a novel series of illusions, where the observer’s action is the key factor. All the observed illusory effects have one feature in common: an illusory impression that the display is “interactive” with the action. We propose that these illusions can be used to understand the neural mechanisms underlying action processing and the perception of sensorimotor interactivity.

### 1.1 The illusions

There are multiple phenomena/demonstrations here, each of which may involve different underlying mechanisms or a combination of them. Here, we will describe each one, along with the procedures and observations.

In all demonstrations, the observer is expected to hold their finger(s) or palm very close to the dynamic random-noise display, but not touch it. They should then execute the following movements, each of which leads to a visual illusion. All these illusions make the visual noise appear interactive with the finger movements.

These illusions can be observed across a wide range of parameters. Dot element size can range from 6 to 95 minutes of visual angle (corresponding to 1 to 16 cell sizes in the video), the frame rate can be 10-60 Hz, and the luminance contrast can range from the highest possible (near 1.0) to just barely detectable, although intermediate to higher contrast is required for stronger effects in most cases. (The readers can directly observe these illusions at: https://sites.google.com/view/magneticsand).

#### 1.1.1 Iconic trace of trajectory

Slowly draw a big trace, such as the letter “S” (Figure 1a), or any other letter. Also, try to draw circles or spirals slowly.

**Figure 1.**
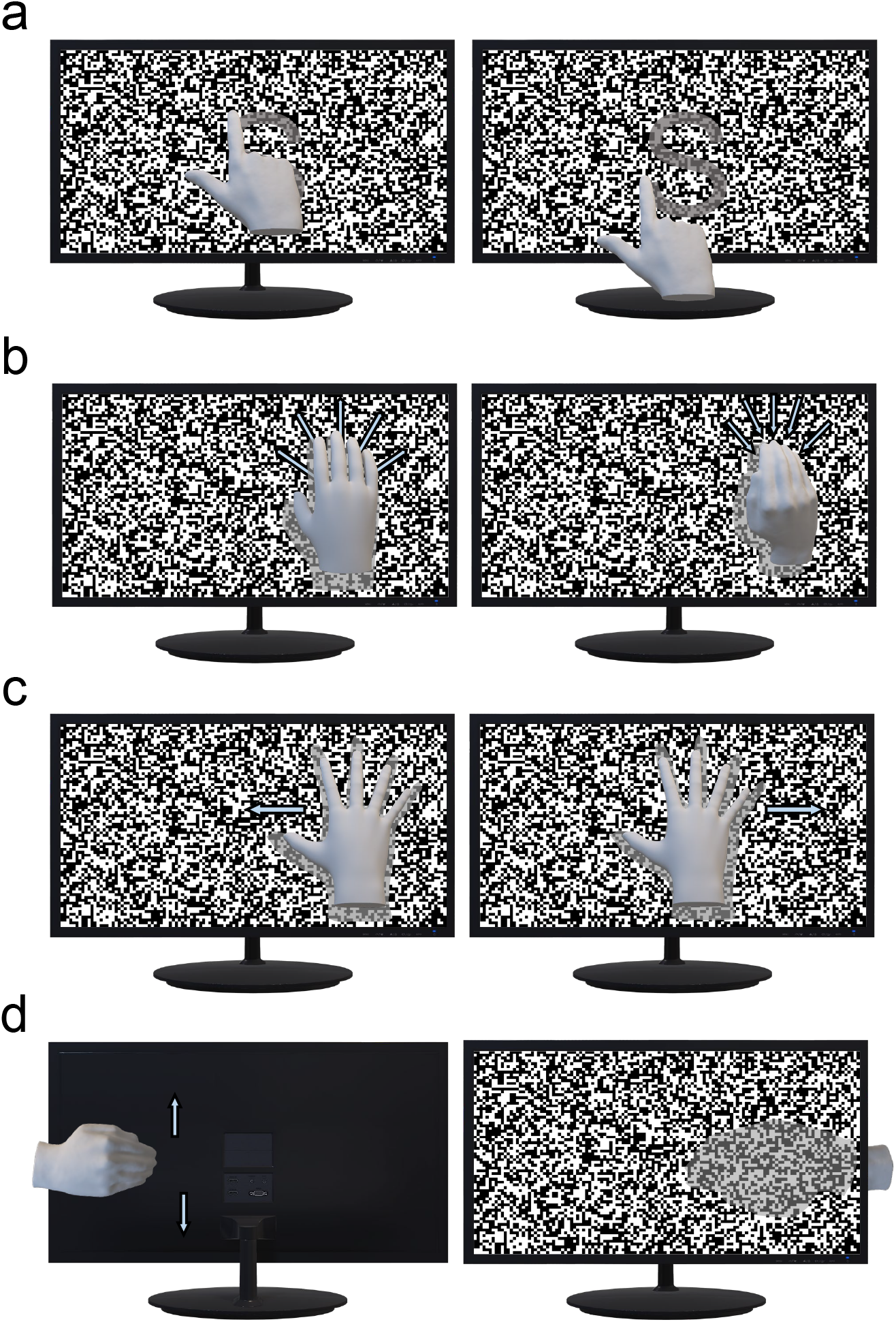
New Illusions. a. **Illusion 1. Iconic Trace of Trajectory**. When the observer draws an “S” letter (or any other letter/pattern) on the dynamic noise display, they see a trace that fades away quickly. b. **Illusion 2. Magnetic Repulsion/Attraction**. Open a palm as it approaches the display, then close the fingers (as though grabbing) as you pull away from it, and repeat. The nearby dots are perceived as though magnetic, moving away while the palm opens and gets close, and being attracted to the fingers when they grab and move away from the display. c. **Illusion 3. Action Capture (with Visible Hand)**. The palm should be wide open, held very close and parallel to the noise display, then slowly moved left and right (or up and down), within a very small spatial range (a slow shudder). It perceptually captures the nearby dots, as though they were being captured and coming together with the palm. d. **Illusion 4. Action Capture (with Invisible Hand)**. While observing the dynamic noise, the observer moves their hand up and down behind the display, so that the hand itself is invisible. The dynamic noise nearby (though often in the spreading area) may appear to be captured by the invisible hand.

> The observer tends to perceive a visible trace of the trajectory (“S” in the example above), which would then appear to decay quickly, in about 0.5 seconds. Some report a brighter trace, some a dimmer shadow along it, etc. One noticeable distinction of this illusion in terms of underlying mechanisms, relative to all the following (illutions 2-4) below, may be that the illusory aspect of the perception itself is not dynamic, but rather static (a trace defined by apparent luminance contrast).

#### 1.1.2 Repulsion/attraction

Slowly open fingers while your palm approaches the display (as though pressing the hand to the surface of the display), then close them while moving away (as though grabbing and picking up sand), as shown in Figure 1b. Repeat. Also, one may try the same with a single finger (as though sticking it to a sticky surface and removing it).

> Observers report both repulsive and attractive “magnetic” interactions with the dots. Dots appear repulsed when the palm gets close to the display and appear attracted to the fingers/palm when the palm moves away with a grabbing motion. Repetition and more focal attention to the hand seems to enhance the effect. Some (though fewer) report that even with a single finger, moving it towards the display (till almost touching) and away yields similar perceptual impressions of the dots escaping away from, and then being attracted towards, the finger.

#### 1.1.3 Action capture (with visible hand)

The palm should be wide open, held very close and parallel to the noise display, then it should slowly move left and right to jitter, with a very small spatial range, as illustrated in Figure 1c. One may also slowly move it up and down, or make a slow big circle. The movement has to be very slow within a small spatial range while keeping the very close distance to the display.

> It seemingly captures the nearby dots, as though they were captured and moving together with the palm.

#### 1.1.4 Action capture (with invisible hand)

While observing the dynamic noise, the observer moves their hand up and down behind the display (so that the hand itself is invisible; see Figure 1d).

> The dynamic noise locally (though often in the spreading area) may appear to be captured by the invisible hand. When the observer pays more attention to their own movement of the hand, the effect tends to get more vigorous.

A closely related action capture can be observed with a very different visual stimulus, i.e., a flickering visual object in the extreme periphery in the temporal frequency range of roughly 0.5-2 Hz. We plan to report more details of this in a separate paper, but it is worth mentioning here because we believe the same or a very similar mechanism underlies the flicker demonstration and this demonstration 1.1.4) above.

## 2 Experiments

Informal observations were gathered by the authors, and in structured experiences during the Demo Night of VSS 2024, with a few hundred audience members.

We performed three psychophysical experiments. In Experiment 1 (Figure 2a). we manipulated cell size and frequency, asking the participants to estimate the magnitude (i.e., strength) of each illusion on a 7-point scale. In Experiment 2, as shown in Figure 2b, we compared static vs. dynamic random noise to see if there are differences across the four illusions. In Experiment 3, as illustrated in Figure 2c, we compared two action conditions: self-action vs. observing others. Magnitude estimation ratings were employed in Experiments 2 and 3 (see more details in the Method section).

**Figure 2.**
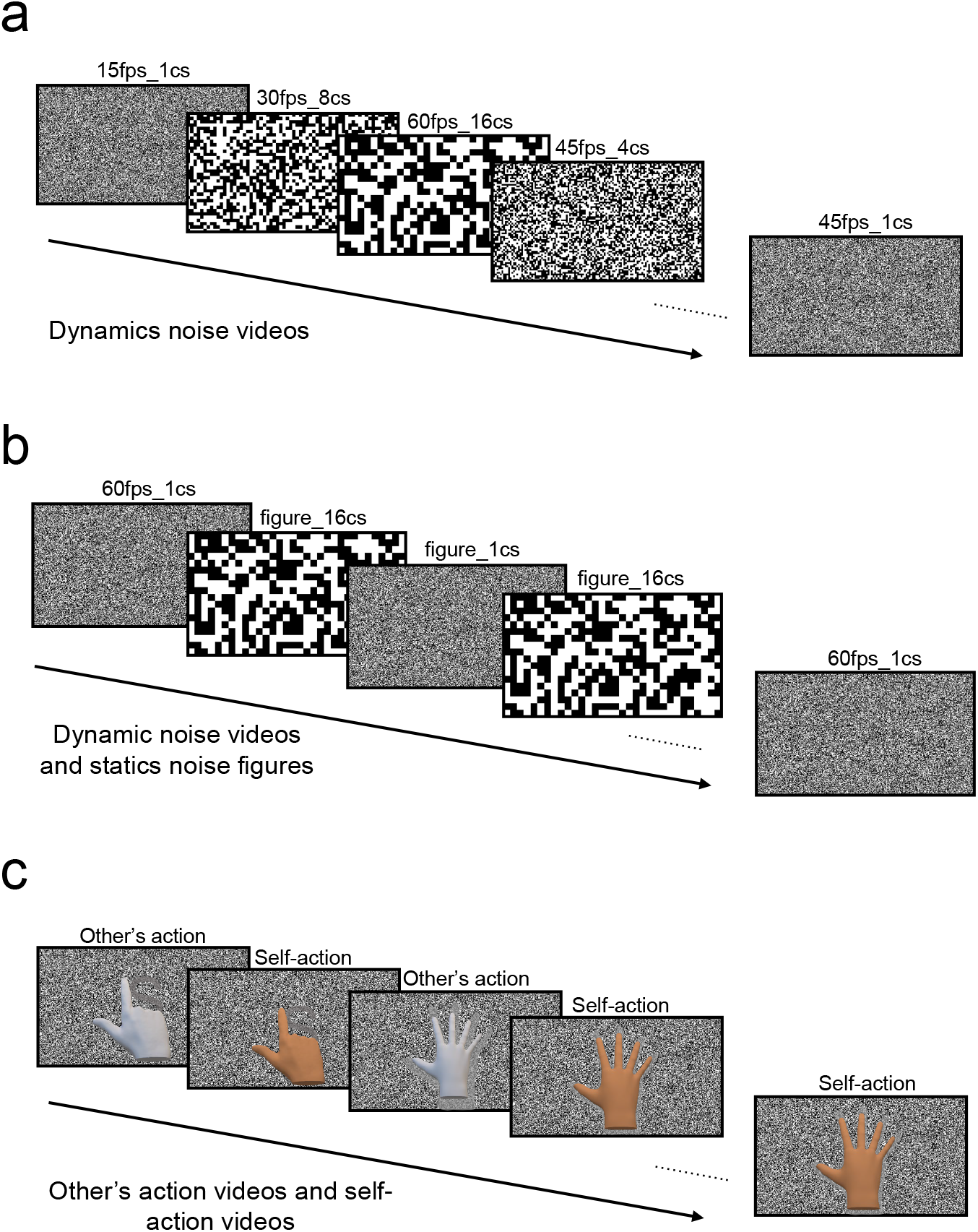
Experimental Procedures. a. Sensitivity to Parameters (Experiment 1: cell size and temporal frequency). b. Comparison of Dynamic vs. Static Noises (Experiment 2). c. Comparison of Self-Action vs. Observing Others (Experiment 3).

The illusions seem to behave somewhat differently in their strength but can be observed within a similar range of parameters, as shown in the results of Experiment 1 (Figure 3a). As shown in the figure, a smaller cell size tends to yield a stronger illusion in general, though it depends on the type of illusion and the participant. Relatively large individual differences were observed regarding which illusion type tends to be the strongest. The low inter-subject correlations across illusions suggest different underlying mechanisms with different weights across observers.

**Figure 3.**
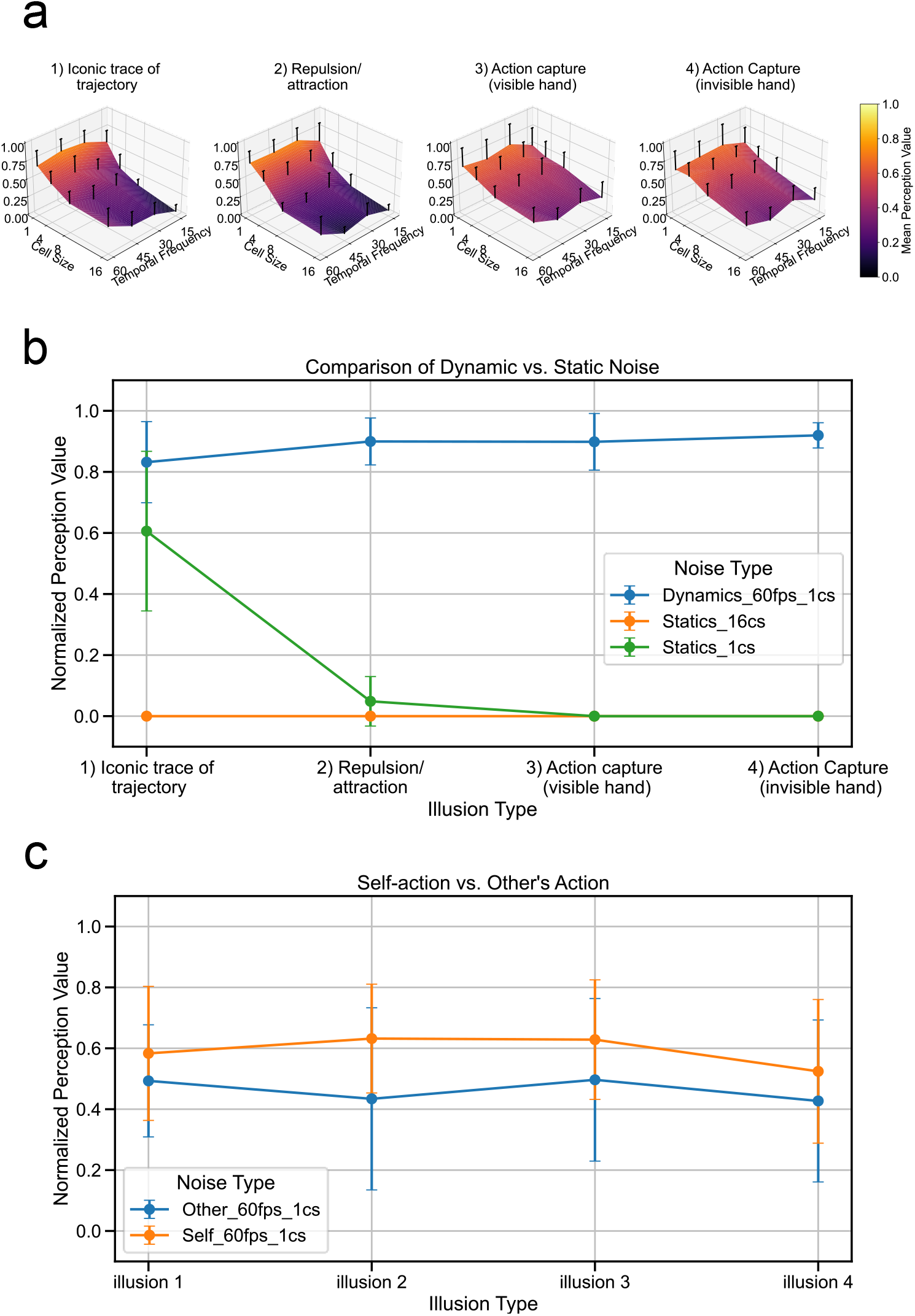
Magnitude Estimation Results in the Three Experiments. a. **Experiment 1: Sensitivity to Parameters**. The horizontal axes represent cell size (1 cell size in the video ≈ 6 minutes of arc) and temporal frequency (fps). The vertical axis shows normalized and averaged magnitude ratings. High frequency and small cell size tend to result in a strong illusion in general. b. **Experiment 2: Static vs. Dynamic Noises**. The X-axis depicts illusion types, while the Y-axis depicts normalized and averaged magnitude ratings. Differently colored curves show results for dynamic noise (60 fps, 1 cs; blue), static noise (16 cs; orange), and static noise (1 cs; green). Illusion 1 (iconic trace) differs significantly from all other types of illusions, as it persists with static noise, especially when the cell size is small (green). c. **Experiment 3: Self-Action vs. Observing Others**. The X-axis depicts illusion types, while the Y-axis depicts normalized and averaged magnitude ratings. Both observation conditions yield illusions, but participants generally report a stronger effect when executing their own actions compared to observing others.

Figure 3b shows the results of the comparison between static vs. dynamic random noise (Experiment 2). As evident from the figure, illusion 1 (iconic trace) behaves very differently from all the other illusions (2-4). While the static noise (regardless of the cell size) shut off all the other three illusions, illusion 1 (iconic trace) seems to be still present, especially when the cell size is small (1 cell size in the video ≈ 6 minutes of arc). This suggests that the iconic trace illusion may have a different underlying mechanism, unique among the four classes of effects.

Figure 3c shows the results of the comparison between self-action vs. observing others’ action (Experiment 3). The first point to note in the results is that both conditions (self-action and observing others’ action) yield robust illusions. The second point is that self-action tends to yield a stronger illusion compared to observing others’ actions across the four types of illusions.

As additional informal observations, several experienced observers compared monocular and binocular observations. The results varied across the effects and the observers, but no unanimous or strong binocular (or monocular) dominance was observed.

## 3 Underlying mechanisms?

We would like to argue that each of the effects described above may necessitate different mechanisms. We describe an intrinsically static adaptation and de-adaptation mechanism which may form the seeds for the first illusion (the iconic trace), and motion-related mechanisms for the rest (illusions 2-4). Beyond these low-level visual effects is the mechanism of action capture, which amplifies the above effects.

1. **(De)adaptation**. The fingers or palm occlude a part of the dynamic-noise display momentarily (though very briefly; perhaps in the order of teens to hundreds of ms), and then slide away. Especially in the case of drawing a letter or a circle, the locally occluded parts undergo a sequence of being occluded, thus quickly de-adapted, and then becoming visible again. This quick refreshment of stimuli may be perceived as a stronger perceptual trace. This explains the seeming brightness or “shadow” in the trace reported by some observers. The exact visual attributes that are refreshed remain an open question. Simple luminance, first-order or second-order luminance contrast [4], and local motion signals would be good candidates.

2. **Accretion/Deletion**. Accretion/deletion is a known motion cue (Figure 4a). It has typically been described when the occluding surface and edge are stationary, but here we propose a special case of dynamic accretion/deletion. Similar to the well-known aperture problem (Figure 4b), each dot in the dynamic noise display presumably has a uniform distribution (with some random noise) across 360 degrees of motion vectors (Figure 5a). Now assume a finger/palm abruptly occludes one side (approximately 180 degrees) of such a vector distribution for the dots along the finger edge (dynamic deletion case; Figure 5b). The vector distribution should now be biased in the general directions moving away from the finger edge. Conversely, when the finger/palm is abruptly removed from a local region, the vector distribution changes from 180 degrees to 360 degrees, thus the occluded half is quickly added (Figure 5c). Together with the de-adaptation effect (mentioned above), it may enhance the perceived direction of the local dots towards the finger/palm. Such effects are consistent with illusions 2 and 3 above.

**Figure 4.**
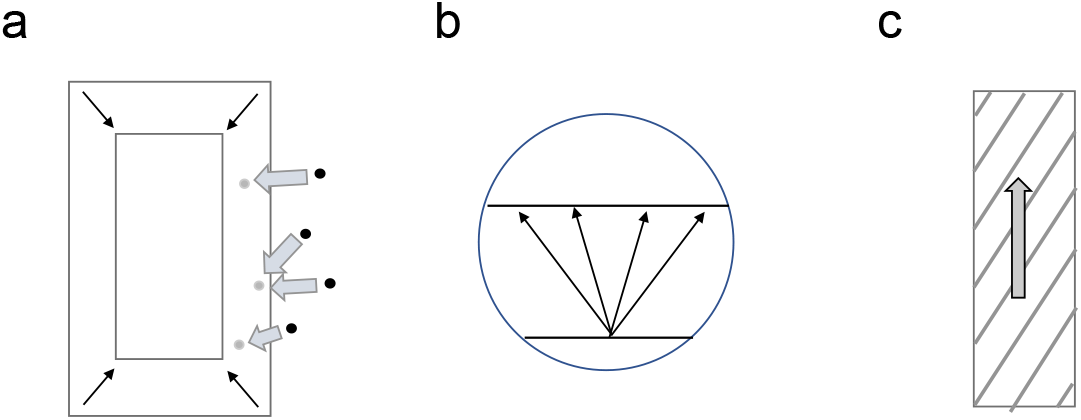
Potential mechanisms underlying the illusions. a. **Accretion/Deletion Cues**. When an occluder (such as a palm or a finger) moves away from the display (as shown by thin oblique arrows), it occludes fewer dots, with some appearing (as shown by the gray dots on the right side; the same occurs on the left side as well). These are perceptually paired with neighboring dot(s), generating new apparent motions (illustrated by gray arrows). When the occluder occludes some nearby dots, the nearby remaining dots may appear to move repulsively away (see more in Figure 3). b. **The Classical Aperture Problem**. When a line translates (from the bottom to the top) within a limited observation window (which can be interpreted as a receptive field of a motion detector), the actual direction of motion can vary across a wide range of angles, as illustrated by the arrows, introducing the concept of visual motion ambiguity. c. **The Barber Pole Illusion**. In the case of the barber pole illusion, the aperture problem (B above) is solved by the unambiguous motion of the terminators (i.e., the crossing points of the stripes with the occluding edges). Thus, the oblique stripes appear to move vertically (or horizontally) in a vertically (or horizontally) elongated window. Our dynamic random dot display differs in the stimulus domain, but a similar ambiguity analysis and solution can be proposed, as described in Figure 3.

**Figure 5.**
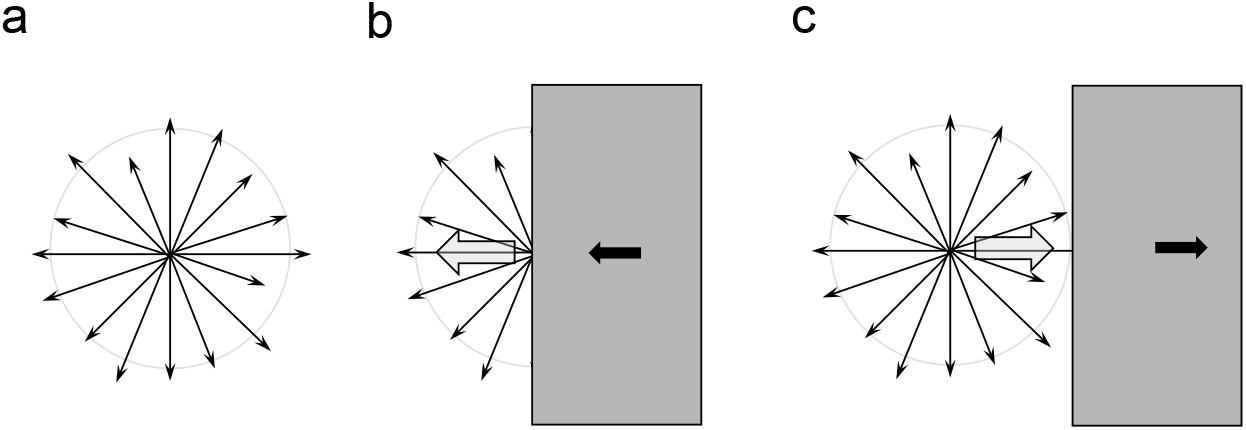
Motion Vector Distribution in the Dynamic Random Noise and How Partial Occlusion Yields Apparent Motion Biases. a. Local motion detectors are stimulated by the dynamic noise in all directions, though with stochastic and varying strength across a 360-degree angle, as illustrated here. b. When an occluder (such as a palm or a finger) moves from right to left, dynamically occluding part of the vector distribution (as shown by the large gray rectangle and the small black arrow in this example), the remaining half of the vectors yield the overall impression of dots moving away repulsively from the occluding edge (thus towards the left), as indicated by the big gray arrow. c. When the occluder moves back from left to right, the left half of the vectors (in this example) reappears, thus inducing an overall impression of the dots moving towards and being attracted to the occluding edge (i.e., right).

The limited availability of such local motion interpretation due to partial occlusion is also known in the case of the barber pole illusion (Figure 4c), where motion-ambiguous oblique stripes appear to be captured by the unambiguous motion of their terminators along the occlusion edges[5]. Indeed, some careful observers, including some of the authors, have reported a very subtle unique percept along the contour of fingers, such as an enhanced contour or a narrow stream line (of dots moving) along the contour of the finger, which could be attributed to the barber pole-like unambiguous motion signals along the contour. However, such an effect is inevitably weak due to the constant movements of the fingers (and still intrinsic ambiguity in the opposing 180-degree directions). When such cues are weak, as in this case, the brain tends to go with population coding, i.e., the grand average of the direction-specific motion detectors’ signals, as demonstrated psychophysically [6, 7] and electrophysiologically [8]. This itself is not explicitly observed in our case of the dynamic random noise, but how it potentially interacts with the other visual motion mechanisms mentioned above is yet to be studied.

3. **Action capture**. The observer’s own active action (even without its own visibility) is known to capture motion-ambiguous stimuli[9], and can even cause organized visual perceptions of motion and form in total darkness[10, 11].In our case, it captures dynamic random stimuli in its direction. Part of our demonstration (3) may be caused by such an active capture mechanism, although the dynamic accretion/deletion may add noise to the effect.

As mentioned above, these mechanisms may contribute differently or with different weights to each illusion, causing individual differences in the relative strength of the illusions. To further reinforce this point, we found that for the first illusion (the iconic trace of trajectory, as shown in Figure 6), even a static random-dot pattern works, but not for all the other illusions (2, 3, and 4). This is highly consistent with our conjecture on the mechanisms: the (potentially static) luminance or contrast adaptation is critically involved in the mechanism underlying the iconic trace illusion, but motion-specific mechanisms are needed for all the other illusions.

**Figure 6.**
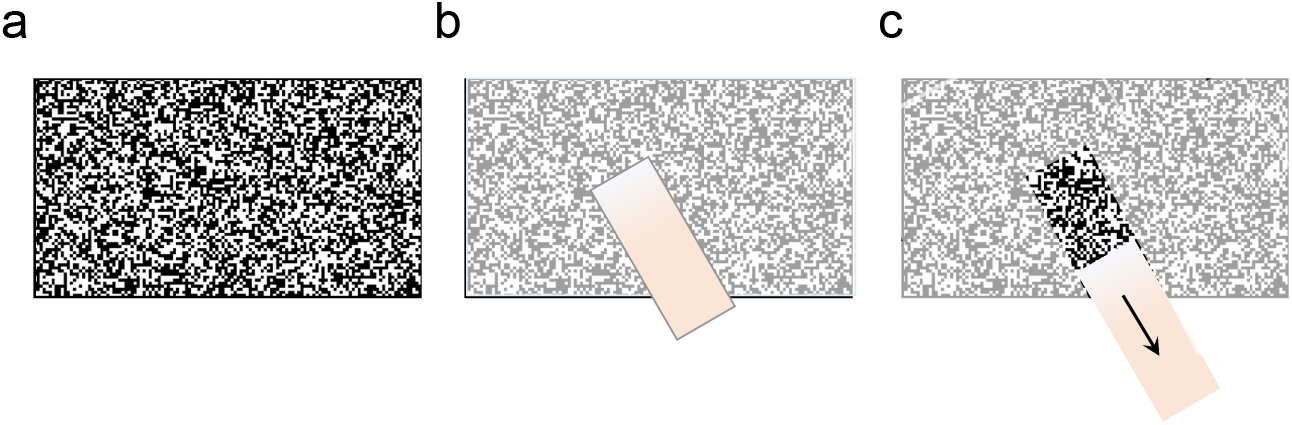
Mechanism underlying iconic trace illusion? a. Original random dot display, undergoing luminance (contrast) adaptation. b. Subjective contrast is attenuated everywhere, except for the finger-occluded region, which escapes from adaptation. c. When the finger moves away, the trace is refreshed, appearing as higher contrast.

What is common and surprising across these illusory effects is that the display appears to change its appearance contingent upon the observer’s hand/finger action nearby, even though the noise display is entirely independent and random. How could this be possible? We would argue for several levels/types of accounts. First, action introduces differences in visual feedback (especially occlusion and de-occlusion in the current study). Second, the brain tunes up or facilitates sensory sensitivity along the action trajectory and direction, presumably to enhance sensory-motor control. Third and more generally, perception and action are coupled for more adaptive behavior in the environment [12]. Thus, in one sense, all the apparent interactivity here is caused by the intrinsically interactive nature of the human sensory systems.

## 4 Method

### 4.1 Participants

Eight participants (7 male, 1 female) took part in the study. All participants provided informed consent prior to their involvement. Other than these laboratory experiment participants, all the effects reported here were tested, improved, and confirmed as demonstrations with 50-70 audience members each year at the Demo Night, the Vision Science Society Annual Meetings in 2019, 2022, 2023, and 2024.

### 4.2 Stimuli

Three types of stimuli were used across all experiments:

1. **Noise Videos**: Created by code, each with a duration of 5 seconds and looped during the experiments, with varying temporal frequencies (15, 30, 45, 60 Hz) and cell sizes (1, 4, 8, 16 pixels).
2. **Noise Figures**: Static images created by code, with varying cell sizes (1, 4, 8, 16 pixels).
3. **Videos of Others’ Actions**: Recorded using a professional camera based on the 24.6MP IMX 410 sensor with a resolution of 1080P, a frame rate of 59.94 frames per second, using the H.264 codec in an MOV (All-I) container, with a bitrate of 240 Mbps. Video processing was conducted using Davinci Resolve to extract clips corresponding to the interactive actions for each illusion. Processed videos maintained a resolution of 1080P, frame rate of 59.94 frames per second, codec H.264, container MP4, and bitrate of 240 Mbps.

### 4.3 Experimental Procedures

#### 4.3.1 Experiment 1: Sensitivity to Parameters

Participants were exposed to noise videos with varying temporal frequencies (15, 30, 45, 60 Hz) and cell sizes (1, 4, 8, 16 pixels), resulting in 16 combinations. Each combination was tested three times, totaling 48 trials (16 parameter combinations × 3 repetitions). Participants performed interactive actions with the noise videos three times per parameter combination, resulting in four interactions per trial. The sequence of trials was randomized.

#### 4.3.2 Experiment 2: Comparison of Dynamic vs. Static Noise

Dynamic noise was represented by a video with a 60 Hz temporal frequency and a 1-pixel cell size. Static noise was represented by images with cell sizes of 1 and 16 pixels.

Participants rated one dynamic video and two static noise images in a randomized sequence, repeated three times, totaling 9 trials (3 conditions × 3 repetitions). Each trial involved four interactions with the dynamic video or static images.

#### 4.3.3 Experiment 3: Comparison of Self Action vs. Observing Others

Participants viewed a recorded video of another participant’s hand interacting with dynamic noise (Other condition) and then mimicked the same action themselves (Self condition) with a dynamic noise video. For each illusion, participants first rated the observed action (Other condition) and then their own action (Self condition). This sequence was repeated for each of the four illusions, with each illusion being tested three times, resulting in 24 trials (4 illusions × 2 conditions (self and other) × 3 repetitions).

### 4.4 Data Collection and Analysis

Participants used a 7-point scale to provide perception values (magnitude estimation ratings) for each illusion. The observation distance for illusions 1 to 3 was approximately 40 to 80 cm, and for illusion 4, it was approximately 20 to 60 cm. Normalized perception values for each illusion were averaged per trial, then across all participants to obtain the mean perception value.

## 5 Data and Code availability

All data and code used in this research are available at the GitHub repository: https://github.com/cantonsir/Magnetic-Sand-Illusions.

## 6 Acknowledgment

The project was supported by SONY Research Award Program. We thank Jiahe Yue for her suggestion to employ a static noise.

## References

[1] Cavanagh, P.: Attention-based motion perception. Science 257(5076), 1563–1565 (1992)

[2] Davidenko, N., Heller, N.H., Cheong, Y., Smith, J.: Persistent illusory apparent motion in sequences of uncorrelated random dots. Journal of Vision 17(3), 19–19 (2017)

[3] Allen, A.K., Jacobs, M.T., Davidenko, N.: Subjective control of polystable illusory apparent motion: Is control possible when the stimulus affords countless motion possibilities? Journal of Vision 22(7), 5–5 (2022)

[4] Lu, Z.-L., Sperling, G.: Second-order illusions: Mach bands, chevreul, and craik-o’brien-cornsweet. Vision Research 36(4), 559–572 (1996)

[5] Vallortigara, G., Bressan, P.: Occlusion and the perception of coherent motion. Vision Research 31(11), 1967–1978 (1991)

[6] Julesz, B.: Foundations of cyclopean perception. (1971)

[7] Braddick, O.: The masking of apparent motion in random-dot patterns. Vision Research 13(2), 355–369 (1973)

[8] Newsome, W.T., Britten, K.H., Movshon, J.A.: Neuronal correlates of a perceptual decision. Nature 341(6237), 52–54 (1989)

[9] Hu, B., Knill, D.C.: Kinesthetic information disambiguates visual motion signals. Current Biology 20(10), 436–437 (2010)

[10] Hofstetter, H.W.: Some observations of phantom visual imagery. Optometry and Vision Science 47(5), 361–366 (1970)

[11] Brosgole, L., Neylon, A.: Kinetic visual imagery. Perceptual and Motor Skills 37(2), 423–425 (1973)

[12] Parr, T., Pezzulo, G., Friston, K.J.: Active Inference: the Free Energy Principle in Mind, Brain, and Behavior. MIT Press, Cambridge, MA (2022)

